# Outcomes of Isoniazid Preventive Therapy among people living with HIV in Kenya: A retrospective study of routine health care data

**DOI:** 10.1101/2020.06.01.127431

**Authors:** Muthoni Karanja, Leonard Kingwara, Polly Kiende, Philip Owiti, Elvis Kirui, Faith Ngari, Elizabeth Onyango, Catherine Ngugi, Maurice Maina, Enos Masini

## Abstract

**Introduction:** Isoniazid preventive therapy (IPT) taken by People Living with HIV (PLHIV) protects against tuberculosis (TB). Despite its recommendation, there is scarcity of data on the uptake of IPT among PLHIV and factors associated with treatment outcomes. We aimed to determine the proportion of PLHIV initiated on IPT, IPT treatment outcomes and screening for TB during and after IPT.

**Methods:** A retrospective cohort study of a representative sample of PLHIV initiated on IPT between July 2015 and June 2018 in Kenya. We abstracted information on socio-demographic, TB screening practices, IPT initiation, follow up, and outcomes from health facilities’ patient record cards, IPT cards and IPT registers. Further, we assessed baseline characteristics as potential correlates of developing TB during and after treatment and IPT completion using multivariable logistic regression.

**Results:** We enrolled 138,442 PLHIV into ART during the study period and initiated 95,431 (68.9%) into IPT. Abstracted files for 4708 patients initiated on IPT, out of which 3891(82.6%) had IPT treatment outcomes documented, 4356(92.5%) had ever been screened for TB at every clinic visit and 4,243(90.1%) had documentation of TB screening on the IPT tool before IPT initiation. 3712(95.4%) of patients with documented IPT treatment outcomes completed their treatment. 42(0.89%) of the abstracted patients developed active TB, 16(38.1%) during and 26(61.9%) after completing IPT. Follow up for TB at 6-month post-IPT completion was done for 2729(73.5%) of patients with IPT treatment outcomes. Sex, Viral load suppression and clinic type were associated with TB development (p<0.05). Levels 4, 5, FBO, and private facilities and IPT prescription practices were associated with IPT completion (p<0.05).

**Conclusion:** Two-thirds of PLHIV were initiated on IPT, with a high completion rate. TB screening practices were better during IPT than after completion. Development of TB during and after IPT emphasises need for keen follow up.

## Introduction

Tuberculosis (TB) is the most prevalent opportunistic infection among People Living with HIV (PLHIV)^1^. It remains the leading cause of death among the HIV population, accounting for one-in-three AIDS-related deaths^1^. Exposure to the *Mycobacterium tuberculosis* bacteria leads to TB infection, which can be controlled and remain inactive for years, but it can also progress to active TB^2^. Overall, the risk of developing TB is more than 20 times greater among PLHIV than those who do not have HIV infection^3^. The risk of TB in HIV-infected persons continues to increase as HIV disease progresses, and immunity decreases^4^. The sustainable development goals (SDGs) and the World Health Organization (WHO) END TB Strategy aim to end the global TB epidemic by 2030^5^.

Kenya is listed by the World Health Organization (WHO) as among the 30 high burden TB states currently facing the triple burden of TB, TB/HIV, and MDR TB with a reported annual incidence of 292 per 100,000 population^6^. Despite the considerable investment done by the government and partners in TB care and prevention in the past 20 years, the disease is still the leading cause of death among PLHIV in the country^7^. According to the Kenya Tuberculosis Prevalence Survey 2016, half of all patients who fall ill to the disease go undiagnosed and untreated^8^. The survey identified HIV as a significant risk factor, contributing to 17% of the overall TB burden^8^. The HIV co-infection rate among notified TB patients in Kenya is at 27%^8^. There is a generalized HIV epidemic in Kenya, but this also varies by geographical areas. Kenya HIV 2018 estimates the HIV prevalence among adults at 4.9% -- Males 3.5%, Females 6.2% with 1,493,400 (1,388,200 adults and 105,200 children) living with HIV^9^. WHO estimates that 13,000 PLHIV died from TB in 2018, a preventable and treatable disease^6^.

In 1993, the WHO issued the first policy statement that recognized the efficacy of management of latent TB infection among PLHIV with isoniazid preventive therapy (IPT). WHO and Stop TB partnership in 2004 then developed and adopted an interim policy on the 3I’s for HIV/TB collaborative activities. The 3Is covers the aspects of intensified TB case-finding and ensuring high-quality anti-tuberculosis treatment, initiation of TB prevention with IPT, and providing control of TB Infection in health-care facilities and congregate settings^10,11^. In 2009, Kenya adopted the WHO 3I’s and two more interventions that included: Immediate ART Therapy and Integration of TB and HIV—collectively dubbed the 5I’s ^5^.

Isoniazid prophylaxis taken for six months offers protection against TB for at least two years^12^. IPT for PLHIV was first recommended by WHO in 1998 and adopted by Kenya for rollout from 2015. After the implementation, the scale-up of IPT among PLHIV initially had a slow start but later peaked in 2016 as derived from National Data Health Information system (**Figure 1**). However, there is limited information on treatment outcomes, including limited published data offering insights into potential gaps and strengths of the implementation of IPT in Kenya. To address this, we carried out this evaluation to assess outcomes of IPT services among PLHIV between July 2015 and June 2018. Specific objectives were: To determine the proportion of PLHIV in care who initiated IPT during this period; b) To assess TB screening practices before IPT initiation; c) To establish IPT treatment outcomes and d) To investigate the proportion that developed TB during the course of treatment and six months post completion. The findings will help inform the national HIV and TB programs and stakeholders on the progress of IPT implementation and help measure progress towards meeting TB control targets.

**Figure 1:**
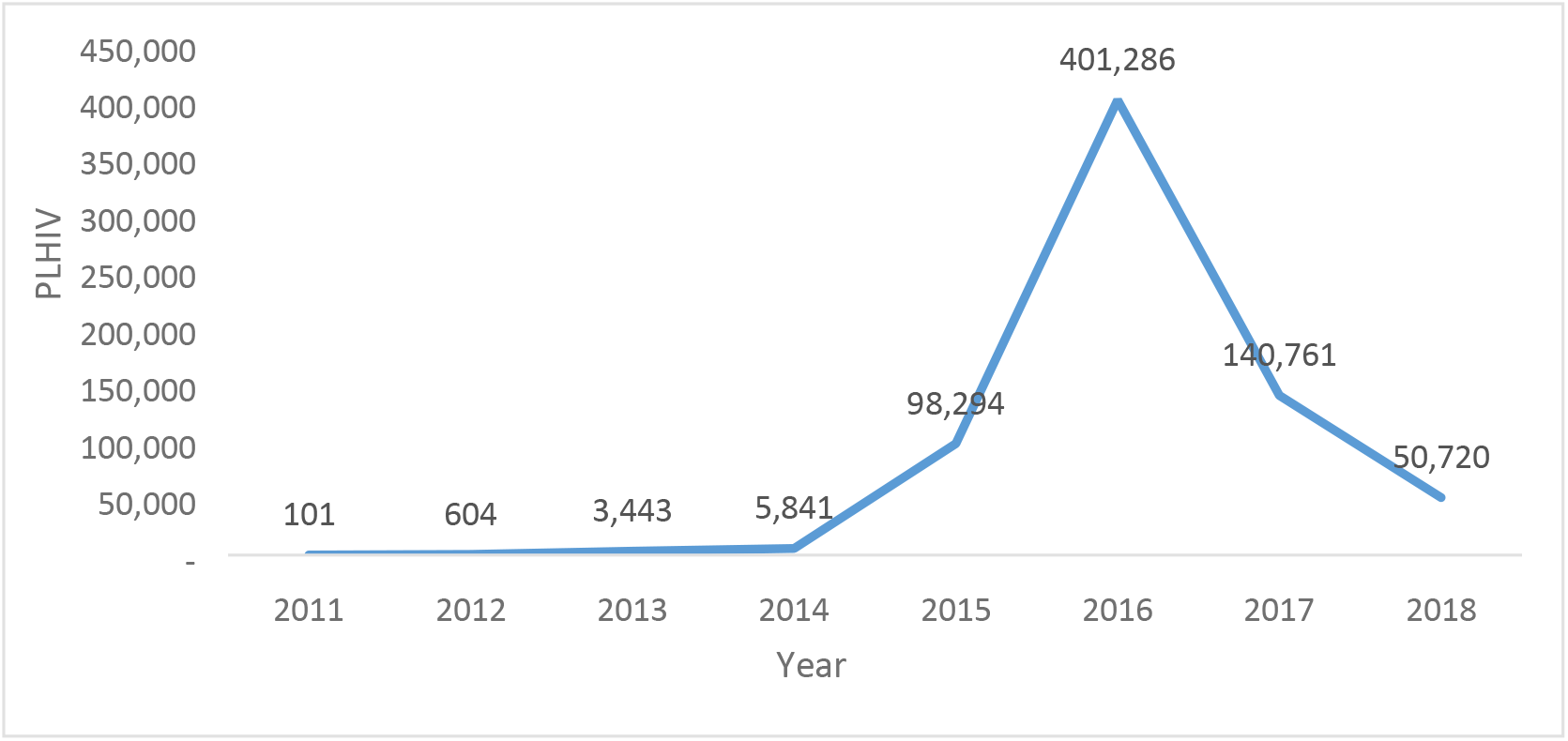
The use of IPT among PLHIV from 2011 to 2018 in Kenya-. The peak in 2015-16 is attributed to both the development of IPT policy by the ministry of health and 100-days rapid results initiative(RRI)

## Material and methods

### Study design

A retrospective study of PLHIV enrolled in care and treatment and initiated on IPT.

### Study setting

We conducted the study in 30 out of the 47 counties of Kenya. In each county, we selected representative facilities from both the urban and the rural setup with a different burden of TB/HIV.

### Study population

All PLHIV in care and initiated on IPT between 1^st^ July 2015 through 30^th^ June 2018.

### Sample size determination

The required sample size was determined using a single population proportion formula n= (z2 * P (1-P)/ d2), and estimations for calculating sample size derived from findings of a study done in Nairobi, Kenya^13^, study precision of 5% and confidence co-efficient of 1.96. We obtained a minimum sample size of (1,700 × 2) + 10% = 3,740.

### Sampling Procedure

We randomly sampled 30 out of 47 counties. Further, the study purposely selected the county referral hospital, a sub-county referral hospital, a private hospital, and a faith-based facility from each of the sampled counties as an adequate representation of facility levels and types. The ultimate sampling unit from the facility was the Comprehensive Care Centers offering IPT services and the patient records within the selected facilities. From each sampling unit, we developed a sampling frame from the IPT patient registers and electronic medical registers where possible. From these registers, we developed a list of eligible patients. The sample size of 3,740 was shared equally among sampled facilities that offer ART services in each county; hence approximately 32 patient files per facility were reviewed. Within the selected facilities, the allocated sample size was divided equally among the three years of the study period. We determined the random sampling interval of patient files by dividing the total number of patients started on IPT that year by the sample size allocated for the same year to get the sampling interval.

### Data collection and management

The study team abstracted aggregate data for patients initiated on IPT and PLHIV on ART for the study period in the selected facilities into MS Excel using a standard patient-specific electronic tool. The tool was pre-tested in a level 4 facility. The data source was from the MOH recording tools in the health facilities, which included patient record card/file, ICF/IPT card (MOH257), and IPT registers. Information on socio-demographic characteristics (age, sex), TB screening practices, IPT initiation, IPT follow up, and IPT outcomes (completed, stopped/discontinued, loss to follow-up, died, and transfer out) were extracted. We also collected data on TB diagnosis during IPT and after IPT completion. Data collection was by trained data collectors under the supervision of study coordinators. The data was backed-up regularly, and confidentiality maintained following national standards.

### Data Analysis

Data was entered, cleaned, and analyzed using SPSS. We then calculated frequencies and proportions for categorical variables and appropriate measures of central tendency and dispersion for continuous variables. Further, we compared patients’ characteristics and clinical practices to IPT completion and other study outcomes using Pearson’s chi-square test and relative risk as appropriate. Factors influencing the primary outcome, IPT completion, were assessed by logistic regression at both bi-variable and multivariable levels and effects presented as odds ratios (adjusted and unadjusted odds ratios) with their 95% confidence intervals. Factors with p<0.25 in the bivariate analysis were considered for multiple regression models unless there was collinearity, sparse, or significant missing data. The level of significance was set at *P*-value <0.05.

### Ethical considerations

We maintained the confidentiality of the patient information collected by using codes for patients’ identifiers and data stored in password-protected computers. The facility in charge granted permission to assess the health facility tools for data abstraction. Ethical clearance was obtained from AMREF Ethical and Scientific Research Committee (*AMREF-ESRC P531/2018*).

## Results

We enrolled 138,442 PLHIV into ART during the study period and initiated 95,431 (68.9%) into IPT as per data retrieved from the National reporting warehouse. A total of 4708 patients files were abstracted, 3891(82.6%) had IPT treatment outcomes documented, while 817 (17.4%) did not have documented treatment outcomes. Of the 3891 initiated on IPT, 3712 (95.4%) completed their treatment, 97 (2.5%) stopped/discontinued treatment, 46 (1.2%) were lost to follow up, 22 (0.6%) transferred out and 14 (0.4%) died, 42 (0.89%) developed active TB with 26 (0.55%) being diagnosed after completing IPT(Fig 2). The characteristics of the participants enrolled (N = 4708) are described in table 1. Of these, 1423 (30.2%) were from level 2 facilities, 2845 (60.4%) were from government-run facilities. Among those enrolled on IPT, 4123 (87.6%) were in the CCC clinics. Females were 3159 (67.1%), 4004 (85.05%) were more than 25 years, and 4525 (96.1%) of the study participants were on ART at the initiation of IPT.

**Table 1.** Characteristics of PLHIV who were initiated on IPT in Kenya, between 1st July 2015 and 30th June 2018

| Variables | Frequency | Percentage (%) |
| --- | --- | --- |
| **Sex** |  |  |
| Female | 3159 | 67.1 |
| Male | 1549 | 32.9 |
| **Age Group** |  |  |
| 0 – 9 | 190 | 4.04 |
| 10 – 14 | 151 | 3.21 |
| 15 – 19 | 109 | 2.32 |
| 20 – 24 | 254 | 5.4 |
| 25+ | 4004 | 85.05 |
| **Patients on ART at Initiation of IPT** |  |  |
| Yes | 4525 | 96.1 |
| No | 183 | 3.9 |
| **Facility Level** |  |  |
| Level 2 | 1423 | 30.23 |
| level 3 | 1723 | 36.6 |
| level 4 | 1402 | 29.78 |
| level 5 | 160 | 3.4 |
| **Facility Ownership** |  |  |
| FBO | 1025 | 21.77 |
| GOK | 2845 | 60.43 |
| Private | 838 | 17.8 |
| **IPT Clinic** |  |  |
| CCC | 4123 | 87.57 |
| Inpatient | 28 | 0.59 |
| MCH/PMTCT | 257 | 5.46 |
| Outpatient clinic | 285 | 6.05 |
| TB clinic | 15 | 0.32 |

ART – antiretroviral therapy; FBO – Faith-based organization; GOK – Government of Kenya; IPT – isoniazid preventive therapy; CCC – comprehensive care clinic; MCH – maternal and child health; PMTCT – prevention of mother to child transmission; TB - tuberculosis

**Fig 2.**
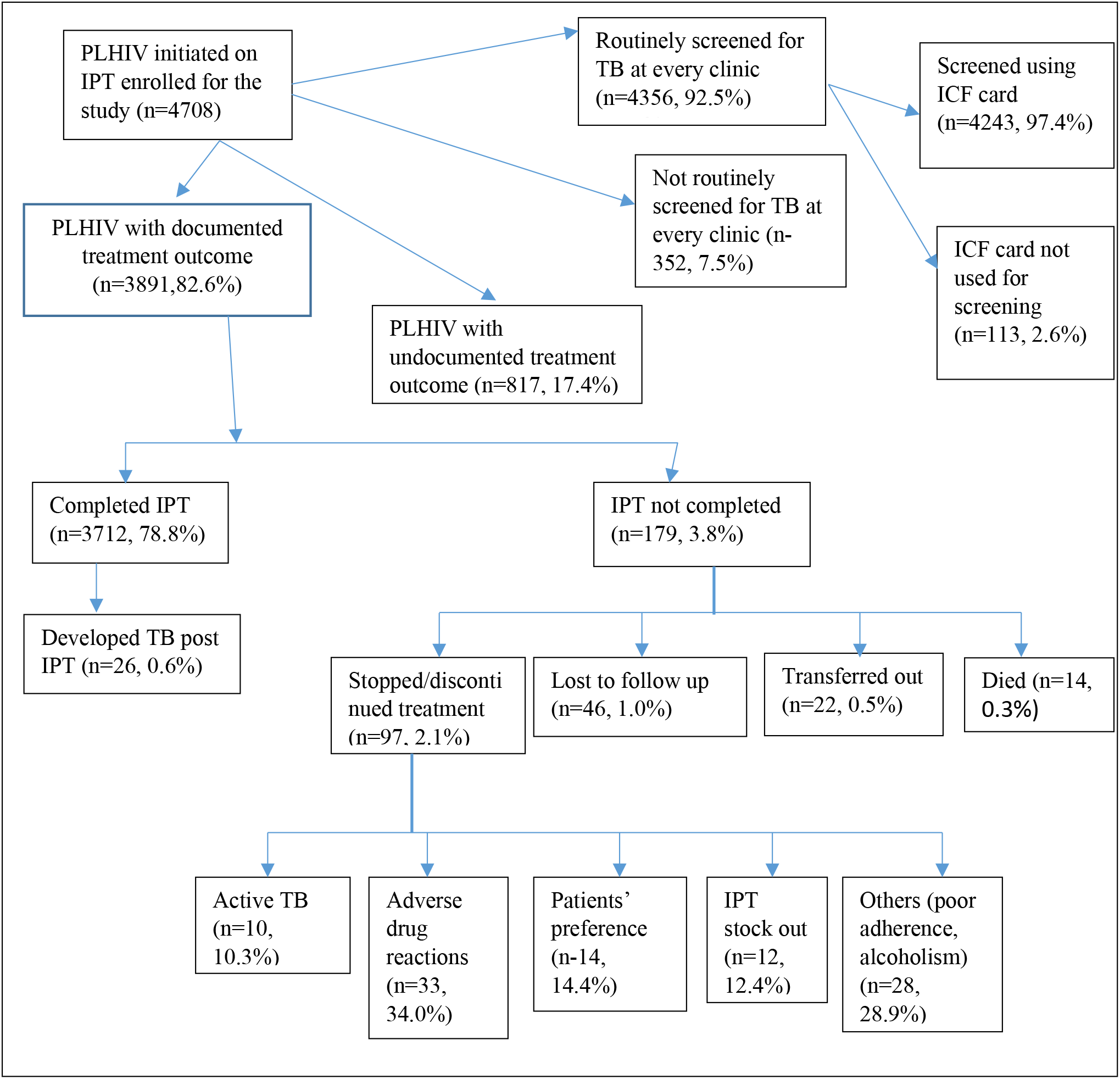
Flow diagram of persons living with HIV who were initiated on Isoniazid Preventive Therapy in Kenya between 1st July 2015 and 30th June 2018.

Those who had ever been screened for TB at every clinic visit were 4356 (92.5%) (Table 2). Among PLHIV screened for TB, 2922 (67.1%) were females, while 3727 (85.6%) were in the age group 25 years and above. Compared to young adults 20-24 years old, the rest of the age groups were more likely to be routinely screened for TB (P<0.05), except for young adolescents 10-14 years old. Of the PLHIV routinely screened, 4,243 (90.1%) had the TB screening documented in the ICF tool before IPT initiation. Among those screened using the ICF tool, 293 (6.9%) were children <15 years, while 3950 (93.1%) were adults (≥15 years). The symptoms assessed in the screening for TB for as documented in the ICF tool for adults included cough (3944, 99.8%), fever (3917, 99.2%), weight loss (3940, 98.7%), and night sweats (3937, 99.7%). TB Symptoms assessed in children were cough (293, 100%), fever (291, 99.3%), weight loss (258, 88.1%), night sweats (251, 85.7%), failure to thrive (190, 64.8%) and lethargy (151, 51.5%).

**Table 2.** Routine TB screening and use of ICF tool for screening of TB among adults and children living with HIV, Kenya, 2015-2018

| Variable | Frequency | Percent (%) |
| --- | --- | --- |
| **Screened for TB routinely (n=4356)** |  |  |
| **Sex** |  |  |
| Female | 2922 | 67.1 |
| Male | 1434 | 32.9 |
| **Age group (years)** |  |  |
| 0 – 9 | 174 | 4.0 |
| 10 – 14 | 137 | 3.1 |
| 15 – 19 | 102 | 2.3 |
| 20 – 24 | 216 | 5.0 |
| 25+ | 3727 | 85.6 |
| **ICF card used** |  |  |
| Yes | 4243 | 97.4 |
| No | 113 | 2.6 |
| **Symptom used for screening** |  |  |
| **Among adults (n = 3950)** |  |  |
| Cough | 3944 | 99.8 |
| Weight loss | 3940 | 99.7 |
| Night Sweat | 3937 | 99.7 |
| Fever | 3917 | 99.2 |
| **Among children (n = 293)** |  |  |
| Cough | 293 | 100 |
| Fever | 291 | 99.3 |
| Weight loss | 258 | 88.1 |
| Night Sweat | 251 | 85.7 |
| Failure to Thrive | 190 | 64.8 |
| Lethargy | 151 | 51.5 |

TB – tuberculosis; ICF – intensified case finding for TB

Of the 42 (0.89%) PLHIV on IPT who developed active TB, 26 (0.55%) were diagnosed after completing IPT while 16 (0.34%) developed TB while on IPT. The median time of being diagnosed with TB after IPT initiation was 2.5 months (interquartile range: 1 - 5 months). Table 3 describes the characteristics of these patients developing TB. Fourteen participants who died while on IPT are described in table 4 - nine were males, median age was 39 years (range 17-65), and none of them had TB diagnosis.

**Table 3:** Characteristics for patients developing TB during and after IPT in Kenya, 2015-2018

TB – tuberculosis; ICF – intensified case finding for TB
| Table 3: Characteristics for patients developing TB during and after IPT in Kenya, 2015- 2018 |  |  |  |  |
| --- | --- | --- | --- | --- |
| Category | Overall<br>n (%) | TB Diagnosis |  | P-value |
|  |  | During IPT<br>n (%) | Post IPT<br>n (%) |  |
| **Age group (yrs)** |  |  |  |  |
| 0 – 9 | 4 (9.5) | 1(6.2) | 3 (11.5) | 0.5 |
| 10 -14. | 1 (2.3) | 0 (0) | 1 (3.8) |  |
| 15 – 19 | 3 (7.1) | 2 (12.5) | 1 (3.8) |  |
| 20 – 25 | 2 (4.7) | 0 (0) | 2 (7.6) |  |
| 25+ | 32 (76.1) | 13 (81.2) | 19 (73.0) |  |
| **Sex** |  |  |  |  |
| Female | 24 (57.1) | 5 (31.2) | 19 (73) | 0.008 |
| Male | 18 (42.8) | 11(68.7) | 7 (26.9) |  |
| **Viral load before IPT initiation** |  |  |  |  |
| Unsuppressed | 6 (14.2) | 1(6.2) | 5 (19.2) | 0.8 |
| Suppressed | 13 (30.9) | 3 (18.7) | 10 (38.4) |  |
| Not Documented | 23 (54.7) | 12 (75) | 11 (42.3) |  |
| **Viral load after IPT Initiation** |  |  |  |  |
| Unsuppressed | 7 (16.6) | 0 (0) | 7 (26.9) | 0.03 |
| Suppressed | 27 (64.2) | 12 (75) | 15 (57.6) |  |
| Not Documented | 8 (19) | 4 (25) | 4 (15.3) |  |
| **Facility type** |  |  |  |  |
| Faith-based | 13 (30.9) | 4 (25) | 9 (34.6) | 0.07 |
| Public | 26 (61.9) | 9 (56.2) | 17 (65.3) |  |
| Private | 3 (7.1) | 3 (18.7) | 0 (0) |  |
| Routine screening |  |  |  |  |
| Yes | 41 (97.6) | 15 (93.7) | 26 (100) | 0.2 |
| No | 1 (2.4) | 1 (6.3) | 0 |  |
| Clinic IPT initiated |  |  |  |  |
| CCC | 39 (92.8) | 13 (81.2) | 26 (100) | 0.02 |
| OPD | 3 (7.2) | 3 (18.8) | 0 |  |

TB – tuberculosis; IPT – isoniazid preventive therapy; CCC – comprehensive care clinics; OPD – outpatient department

**Table 4:**
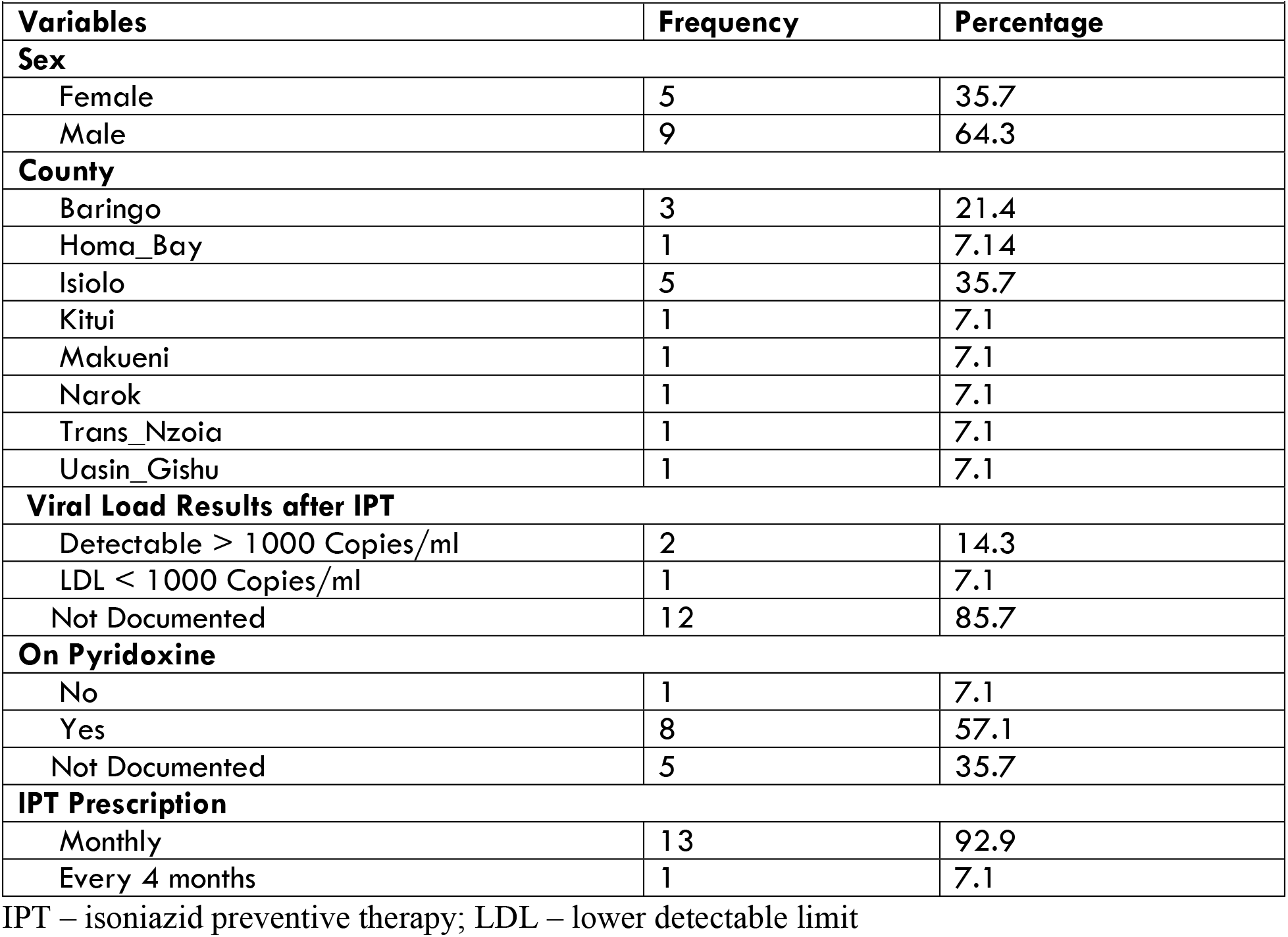
Characteristics for patients who died during IPT in Kenya, 2015-2018 (n=14)

Of the 3712 patients who completed IPT, 2729 (73.5%) were followed up for TB status at six months post-IPT completion. The patient follow-up post-IPT became less frequent after that with 554 (14.9%) followed up at 12 months, 144 (3.9%) at 18 months, and 285 (7.7%) at 24 months as shown in figure 3.

**Figure 3:**
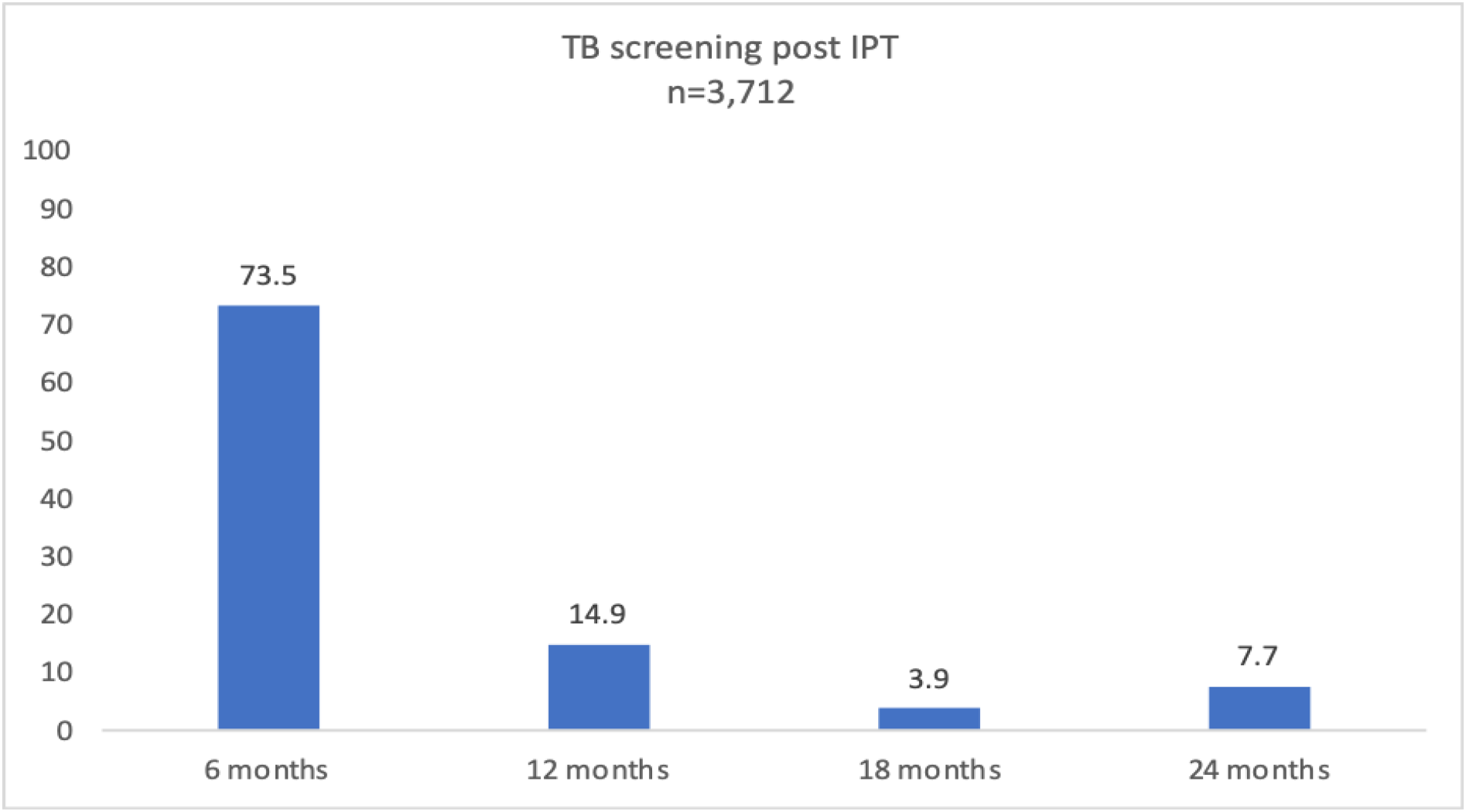
Frequency of TB screening post IPT completion among PLHIV, Kenya, 2015-2018.

Factors significantly associated with completion of IPT included health facility level 4 (aOR 0.50, CI 0.33–0.74; *P* = 0.001), level 5 (aOR 0.39, CI 0.17–0.90; *P* = 0.03); privately-owned facilities (aOR 0.61, CI 0.43–0.87; *P* = 0.006); faith-based organizations (aOR 1.65, CI 1.02–2.68, P=0.04) and IPT prescription practices (monthly, 2 months, 3 months and above 4 months) (Table 5).

**Table 5:**
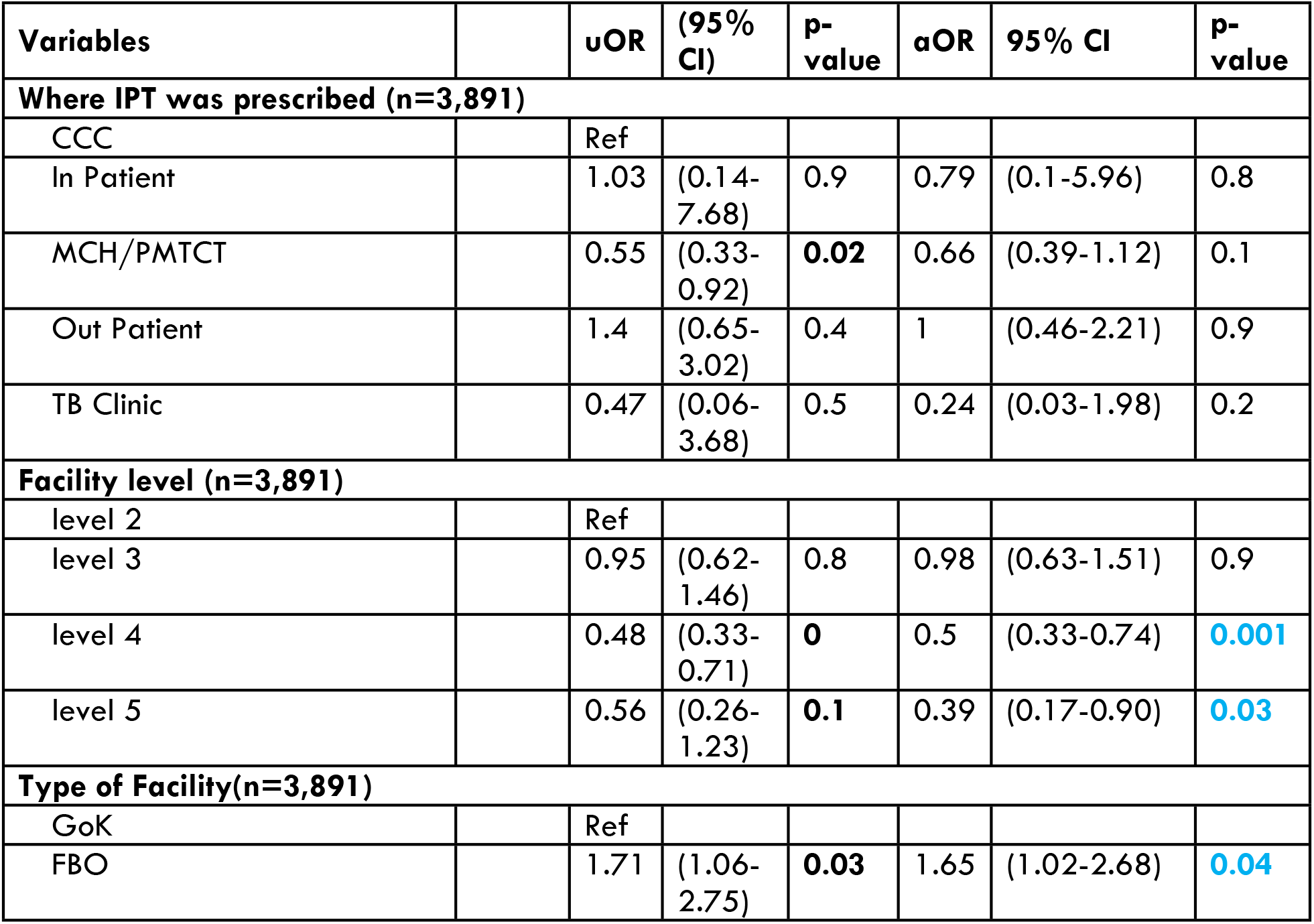
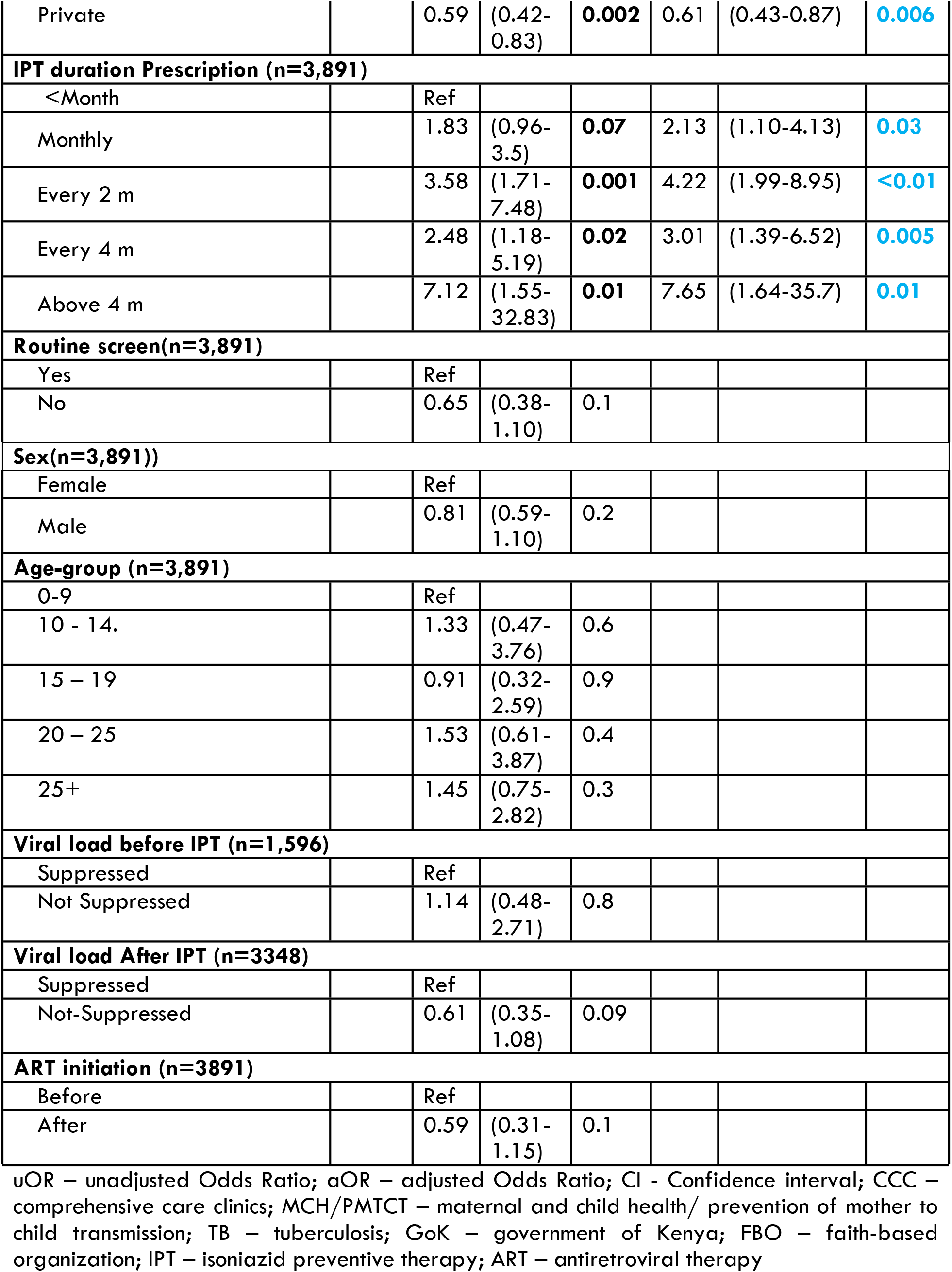
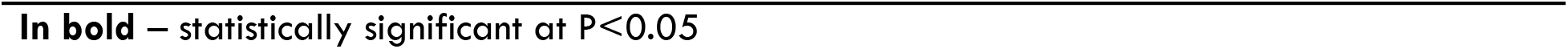
Factors associated with IPT completion among PLHIV in Kenya, 2015-2018

| Variables |  | uOR | (95% CI) | p-value | aOR | 95% CI | p-value |
| --- | --- | --- | --- | --- | --- | --- | --- |
| **Where IPT was prescribed (n=3,891)** |  |  |  |  |  |  |  |
| CCC |  | Ref |  |  |  |  |  |
| In Patient |  | 1.03 | (0.14-7.68) | 0.9 | 0.79 | (0.1-5.96) | 0.8 |
| MCH/PMTCT |  | 0.55 | (0.33-0.92) | **0.02** | 0.66 | (0.39-1.12) | 0.1 |
| Out Patient |  | 1.4 | (0.65-3.02) | 0.4 | 1 | (0.46-2.21) | 0.9 |
| TB Clinic |  | 0.47 | (0.06-3.68) | 0.5 | 0.24 | (0.03-1.98) | 0.2 |
| **Facility level (n=3,891)** |  |  |  |  |  |  |  |
| level 2 |  | Ref |  |  |  |  |  |
| level 3 |  | 0.95 | (0.62-1.46) | 0.8 | 0.98 | (0.63-1.51) | 0.9 |
| level 4 |  | 0.48 | (0.33-0.71) | **0** | 0.5 | (0.33-0.74) | **0.001** |
| level 5 |  | 0.56 | (0.26-1.23) | **0.1** | 0.39 | (0.17-0.90) | **0.03** |
| **Type of Facility(n=3,891)** |  |  |  |  |  |  |  |
| GoK |  | Ref |  |  |  |  |  |
| FBO |  | 1.71 | (1.06-2.75) | **0.03** | 1.65 | (1.02-2.68) | **0.04** |
| Private |  | 0.59 | (0.42-0.83) | **0.002** | 0.61 | (0.43-0.87) | **0.006** |
| **IPT duration Prescription (n=3,891)** |  |  |  |  |  |  |  |
| <Month |  | Ref |  |  |  |  |  |
| Monthly |  | 1.83 | (0.96-3.5) | **0.07** | 2.13 | (1.10-4.13) | **0.03** |
| Every 2 m |  | 3.58 | (1.71-7.48) | **0.001** | 4.22 | (1.99-8.95) | **<0.01** |
| Every 4 m |  | 2.48 | (1.18-5.19) | **0.02** | 3.01 | (1.39-6.52) | **0.005** |
| Above 4 m |  | 7.12 | (1.55-32.83) | **0.01** | 7.65 | (1.64-35.7) | **0.01** |
| **Routine screen(n=3,891)** |  |  |  |  |  |  |  |
| Yes |  | Ref |  |  |  |  |  |
| No |  | 0.65 | (0.38-1.10) | 0.1 |  |  |  |
| **Sex(n=3,891))** |  |  |  |  |  |  |  |
| Female |  | Ref |  |  |  |  |  |
| Male |  | 0.81 | (0.59-1.10) | 0.2 |  |  |  |
| **Age-group (n=3,891)** |  |  |  |  |  |  |  |
| 0-9 |  | Ref |  |  |  |  |  |
| 10 - 14. |  | 1.33 | (0.47-3.76) | 0.6 |  |  |  |
| 15 – 19 |  | 0.91 | (0.32-2.59) | 0.9 |  |  |  |
| 20 – 25 |  | 1.53 | (0.61-3.87) | 0.4 |  |  |  |
| 25+ |  | 1.45 | (0.75-2.82) | 0.3 |  |  |  |
| **Viral load before IPT (n=1,596)** |  |  |  |  |  |  |  |
| Suppressed |  | Ref |  |  |  |  |  |
| Not Suppressed |  | 1.14 | (0.48-2.71) | 0.8 |  |  |  |
| **Viral load After IPT (n=3348)** |  |  |  |  |  |  |  |
| Suppressed |  | Ref |  |  |  |  |  |
| Not-Suppressed |  | 0.61 | (0.35-1.08) | 0.09 |  |  |  |
| **ART initiation (n=3891)** |  |  |  |  |  |  |  |
| Before |  | Ref |  |  |  |  |  |
| After |  | 0.59 | (0.31-1.15) | 0.1 |  |  |  |

uOR – unadjusted Odds Ratio; aOR – adjusted Odds Ratio; CI - Confidence interval; CCC – comprehensive care clinics; MCH/PMTCT – maternal and child health/ prevention of mother to child transmission; TB – tuberculosis; GoK – government of Kenya; FBO – faith-based organization; IPT – isoniazid preventive therapy; ART – antiretroviral therapy
**In bold** – statistically significant at $P < 0.05$

IPT – Isoniazid Preventive Therapy; PLHIV – People Living with HIV; ART – Antiretroviral Therapy; TB – Tuberculosis; ICF – intensified case finding

## Discussion

About two-thirds of the patients initiated on IPT for the period under review were similar to other studies in Africa from 2014–2017 ^14–16^. We did not probe into why the rest were not on IPT, though based on unpublished programmatic data, this can be attributed to being ineligible due to contraindications for IPT: active TB at the time of ART initiation or previous IPT course. The coverage of IPT in 15 high TB/HIV burden countries that reported IPT data to WHO in 2017 ranged from 1% in Eswatini to 15% in South Africa among newly enrolled PLHIV, while 59 countries averaged 36%^17^. Kenya’s comparable data could not be computed for the same period due to differing indicator definitions. However, a study in an urban health center in the capital Nairobi reported a 77% IPT uptake ^13^ amongst adults PLHIV who had been in care for at least six months and regardless of ART status. Though our study focussed on only those on ART, we do not think this differs much as the country’s policy has been initiation of IPT regardless of ART status.

Clinicians routinely used the ICF tool to screen for TB among PLHIV before initiating IPT. Most PLHIV were regularly screened for the four ICF symptoms as recommended. The results confirmed a high coverage of routine screening for TB among most of the PLHIV. However, a few of the PLHIV did not have routine TB screening, and this could be due to poor documentation, as observed in most of the records. Comparable to a previous study in western Kenya, we found much higher screening coverage. This can partly be explained by the increased focus on quality of care interventions among PLHIV in the past 2 years^18^. These findings revealed a better uptake of screening for TB, unlike in the UK, where a clinical audit found that less than a quarter of the PLHIV routinely screened for TB ^19^. The WHO recommends routine TB screening for PLHIV at every clinic visit ^20^. This recommendation has also been adopted as policy in Kenya as stipulated in the National ARV Guideline, 2018 edition^21^ that states, all PLHIVs should be screened for TB at every clinic visit or encounter. Screening PLHIV actively for TB at every clinical encounter allows for provision of TB preventive therapy services among those eligible ^22^. Also, routine screening for TB was applied equally among males and females, but this differed among the age groups with the young adults aged 20–24 years having lower chances of being routinely screened when compared to the rest. The reason for these findings is not apparent but are similar to those reported in a study in western Kenya ^18^. In children, the additional symptoms required for screening, such as failure to thrive, and lethargy were poorly documented, showing missed opportunities for actively looking for TB among them.

The IPT completion rates in Kenya were above 95%, which is higher compared to other African countries, where it ranges from 75–94%^14,16,22,23^. The excellent results are attributable to the adoption and implementation of policy guidelines on IPT in Kenya, integration of tuberculosis and HIV services ^21,24^, sensitization of the health care workers on the importance of IPT among PLHIV and mechanisms to support treatment adherence in HIV care clinics. In Kenya, adverse drug reactions were the leading cause of IPT non-completion. South Africa and Zimbabwe reported similar findings where 3.8% and 7%, respectively discontinued IPT due to side effects ^25^.

Age was not associated with IPT completion rates. A study in Tanzania showed age as a factor with a lower adherence level among patients aged 18-29, 30-49, and ≥ 50 years^26^. A study in Congo showed that higher age at IPT initiation was associated with IPT completion ^27^. Similar to the findings of this study, others have shown that sex, occupation, socio-economic status, duration of HIV infection, being on ARVs, and duration of ARV use were not associated with adherence ^26^.

A study in Swaziland found high IPT completion rates despite the level of facility delivering it, unlike in Kenya, where level 4 and 5 facilities offering IPT were associated with lower treatment completion rates ^22^. The study results showed that IPT clients from private health facilities had lower completion rates. The need for enhanced adherence counselling and active tracing amongst clients on IPT has proved to be vital for successful completion of IPT ^23^. In other studies, clients with HIV WHO stage 3 & 4 and lower CD4 counts were associated with lower completion rates ^28^. Another study showed that participants on ART at IPT initiation were more likely to complete IPT than those who were not ^27^.

The study findings, particularly those on TB as an outcome of those on IPT is lower than that of a similar study conducted among children within an HIV setting in Kenya, where 3% of children initiated on IPT developed TB ^29^. A large retrospective study in Myanmar involving 3375 participants, however, indicated up to (1.2%) of those initiated on IPT were diagnosed with TB, including 0.3% while on IPT and 0.9% within one year of IPT completion. The findings among those who developed TB while on IPT in our study are comparable to the Myanmar study; however, those who developed TB post-IPT was lower (0.6%) compared to the same study (0.9%) ^30^.

Our findings showed that TB screening guidelines were followed up well during the administration of IPT, but after completion of IPT, there was laxity. Only about two thirds of the patients who completed IPT were followed up for TB status at six months post-IPT completion. The patient follow-up post-IPT became less frequent after that with less than a quarter followed up at subsequent months. According to national guidelines, follow up of PLHIV on IPT should be conducted monthly during IPT with rescreening for TB at every visit to help address the adverse drug effects and to detect any signs of active TB for initiation of early treatment. The guidelines ensure patients who have active TB disease do not end up developing drug resistance TB later in life. Also, the patients should be followed up for the next two years post IPT to ensure active TB is detected early.

### Study Limitation

Programmatic data has inadequate data on TB screening for patients who only come for prescription refills, and symptom screening conducted as part of routine clinical services are not available for verification. Additionally, under these normal practice conditions, we were unable to obtain complete data on all patients. Due to the low numbers of PLHIV on IPT who developed TB, it was not possible to establish the relationship between patient characteristics and TB as an outcome. Further, over 60% of patients who developed TB did not have a viral load, hence this could not be evaluated during the characterization.

## Conclusion

Kenya MoH initiated two-thirds of PLHIV on IPT during the study period, and the completion rate was very high. TB screening practices were not up to the standards for all the PLHIV, and a few patients were still diagnosed with active TB during IPT uptake and also post-IPT completion. There remains a gap in TB screening for the PLHIV, and among those screened, the ICF tool is not uniformly applied. IPT completion rates among HIV infected patients were demonstrated to be high. Routine TB screening while on IPT was better than after IPT completion.

There is a need to strengthen TB/HIV integration to ensure all PLHIV are routinely screened for TB and sensitize the health care workers on the use of the ICF tool for screening. More emphasis is required on documentation for IPT clients to reduce the proportion of clients with no treatment outcomes recorded. Quality of care across all health facilities should be enhanced to ensure similar treatment outcomes not based on the level of ownership. The study recommends further evaluation of factors associated with the development of TB during and after IPT completion. Also, we need to assess the quality of screening of TB among PLHIV and conduct routine data audits on ICF.

## Acknowledgment

This 2019 report findings on the assessment of the outcomes of isoniazid preventive therapy among people living with HIV in Kenya is through collaborative efforts of individuals and institutions led by Ministry of Health through the Division of National AIDS and STI Control Program (NASCOP) and Division of National Tuberculosis, Leprosy and Lung Disease Program (NTLD-P). We thank the study co-ordinator from MOH Moseti Makori and the IPT study team members that consisted of a team from Division of NASCOP (Steve Ambune and Evans Imbuki), Division of NLTD-P (Kiogora Gatimbu, Newton Omale, Richard Kiplimo, and Martin Githiomi), MOH (Kigen Bartilol, Maureen Kamene, George Githuka, Stephen Muleshe), NPHL (Josephine Wahogo), KEMRI (Jane Ong’ang’o), KNH (Margaret Oluka), CHS (Lorraine Mugambi-Nyaboga, Evelyne Ng`ang`a and Wandia Ikua), WHO (Hillary Kipruto) and the Global CHAI Team. Special appreciation goes to Global Fund for AIDS, TB and Malaria for the financial support in the implementation of the assessment.

## Author Contributions

MK conceived the study, collected data, analyzed the data and drafted the manuscript; EM supervised data collection, contributed to data analysis and assisted in drafting and submission of the manuscript. LK, PK, EK, FN, PO, MM, EO, CN and EM participated in data collection, data curation, formal analysis, methodology development and review of the manuscript. EK and LK further contributed to data analysis, drafting and critical revision of the manuscript. All authors approved the final version of the manuscript.

